# Comparison of *Yersinia enterocolitica* DNA methylation at ambient and host temperatures

**DOI:** 10.1101/2019.12.16.878991

**Authors:** Dustin J. Van Hofwegen, Carolyn J. Hovde, Scott A. Minnich

## Abstract

Pathogenic bacteria recognize environmental cues to vary gene expression for host adaptation. Moving from ambient to host temperature, *Yersinia enterocolitica* responds by immediately repressing flagella synthesis and inducing the virulence plasmid (pYV)-encoded type III secretion system. In contrast, shifting from host to ambient temperature requires 2.5 generations to restore motility suggesting a link to the cell cycle. We hypothesized that differential DNA methylation contributes to temperature-regulated gene expression. We tested this hypothesis by comparing single-molecule real-time (SMRT) sequencing of *Y. enterocolitica* DNA from cells growing exponentially at 22°C and 37°C. The inter-pulse duration ratio rather than the traditional QV scoring was the kinetic metric to compare DNA from cells grown at each temperature. All 565 YenI restriction sites were fully methylated at both temperatures. Among the 27,118 DNA adenine methylase (Dam) sites, 42 had differential methylation patterns while 17 remained unmethylated regardless of temperature. A subset of the differentially methylated Dam sites localized to promoter regions of predicted regulatory genes including LysR-type and PadR-like transcriptional regulators, and a cyclic-di-GMP phosphodiesterase. The unmethylated Dam sites localized with a bias to the replication terminus, suggesting they were protected from Dam methylase. No cytosine methylation was detected at Dcm sites.

**DATA SUMMARY:** All methylation/base modification data are available at figshare at https://dx.doi.org/10.6084/m9.figshare.3493247 and https://dx.doi.org/10.6084/m9.figshare.3493310.

**IMPACT STATEMENT:** Organisms sense and respond to their environment, in part, by epigenetic variation mediated by DNA methylation. Pathogenic bacteria vary gene expression to allow survival and activate virulence systems in response to host temperature. *Yersinia enterocolitica,* a facultative intracellular pathogen, respond by immediately repressing flagella synthesis and inducing the virulence plasmid-encoded type III secretion system. In this work, we examined the locations of DNA methylation throughout the *Y. enterocolitica* genome. While most methylation target sites were fully methylated, we identified sites with disparate temperature-dependent methylation. Several of these sites were within promoter regions of predicted regulatory genes. Differences in DNA methylation in promoter sequences are often responsible for variations in transcription. Identification of these differences in methylation provide likely candidates for regulators responsible for temperature-dependent phenotypes.

## INTRODUCTION

The adaptation of facultative bacterial pathogens to their mammalian host environment requires significant global changes in gene regulation to establish infection. Host cues utilized for this transition vary among pathogens, but for many bacteria, host temperature is a key environmental signal (Maurelli, 1989). Temperature sensing is especially prominent in the pathogenic *Yersinia. Y. enterocolitica,* a Gram-negative enteropathogen, shows significant phenotypic changes between the narrow range of 30°C and 37°C (host temperature). Changes include a temperature-dependent requirement for calcium ion (2.5 mM at 37°C) (Michiels & Cornelis, 1991; Straley *et al.*, 1993), modification of LPS acylation to circumvent Toll-like receptor (TLR) 4 stimulation (Montminy *et al.*, 2006; Rebeil *et al.*, 2004), metabolic differences such as urease and acetoin production (de Koning-Ward & Robins-Browne, 1997), and the reciprocal temperature-controlled regulation between two type-III secretion systems (TTSS). The latter includes the immediate repression of flagellum biosynthesis and concomitant induction of the pYV (virulence plasmid) TTSS at 37°C (Minnich & Rohde, 2007; Rohde *et al.*, 1994). The reciprocal temperature regulation of these two TTSS may be required because of the substrate reciprocity of their exported proteins (Minnich & Rohde, 2007). Co-expression of flagella and the virulence TTSS at 37°C would result in injection of flagellin into host cells by the pYV-encoded TTSS. Flagellins are potent cytokine inducers of TLRs 5 and Ipaf and their injection into host cells could effectively countermand the pYV-TTSS effectors, termed *Yersinia* outer proteins (Yops). The Yops collectively act to suppress the host innate immune system. Conversely, the flagellar TTSS exports Yops into the host extracellular milieu, rather than direct injection into host cells, diluting their effect. Thus, the immediate cessation of flagellin expression and concomitant induction of the Yops may be essential during the initial stages of host infection.

In contrast to the *Y. enterocolitica* rapid response to host temperature, acclimation to 25°C after a temperature downshift (37°C to 25°C) is much slower. *Y. enterocolitica* adapted to 37°C and downshifted to 25°C requires ~2.5 generations before flagellins *(fleABC*) are expressed (Rohde *et al.*, 1994). The relative timing varies between four and 10 hrs, depending on growth rate (rich vs minimal medium), but the 2.5 generation requirement is consistent. This suggests restoration of a low temperature phenotype is cell-cycle dependent. A model of temperature-dependent differential DNA methylation could link temperature-regulated genes to the cell cycle if expression of a key regulatory gene was sensitive to DNA methylation. This is because two generations of DNA replication are required for a DNA site to go from a fully methylated to an unmethylated state.

Bacterial DNA methylation occurs at adenine and cytosine bases providing epigenetic information in the form of N6-methyladenine (N6mA), N4-methylcytosine (N4mC), and 5-methylcytosine (5mC). DNA methylation is a mechanism to discriminate self from non-self (restriction sensitive), direct mismatch repair, excise non-methylated, i.e. non-self, strands of DNA (Wilson, 1991), and activate transpositions (Roberts *et al.*, 1985). Recent studies expand the role of bacterial DNA methylation to include regulation of cell cycle progression (Collier, 2009; Collier *et al.*, 2007) and modulation of gene expression (Casadesús & Low, 2006; Low *et al.*, 2001; Srikhanta *et al.*, 2010). For example, DNA replication forks leave a wake of transient hemi-methylated sites on the newly synthesized DNA strand. Hemi-methylation of gene promoters can either activate or repress expression by promoting or inhibiting the binding of specific transcription factors. Transient hemi-methylation states associated with the passing of DNA replication forks, account for the link between *Caulobacter crescentus* developmental gene regulation and its cell cycle (Collier *et al.*, 2007) and the frequency of *E. coli Tn10* transposition (Campbell & Kleckner, 1988). Thus, modulation of DNA methylation provides a nondestructive and reversible means of DNA modification. This biphasic epigenetic switch is one mechanism organisms use to sense and adapt to their environment (Atack *et al.*, 2015).

Temperature modulations of DNA supercoiling, histone-like protein DNA binding, and intrinsic DNA bends also contribute to *Y. enterocolitica* host adaptation. DNA supercoiling levels naturally respond to changes in temperature to control expression of flagella and virulence factors or can be artificially manipulated using gyrase inhibitors or novobiocin-resistant mutants (Rohde *et al.*, 1994). Intrinsic DNA bends associated with poly-A and T tracts are sensitive to temperature and effectively melt at 37°C. Changes in DNA structure affect binding of histone-like proteins, promoter function, and methylation (Dorman, 1991). Coupling temperature-regulation to DNA methylation is also evident in *Yersinia.* Overproduction of DNA adenine methylase (Dam) in *Y. pseudotuberculosis* overrides the low temperature repression of Yop expression, but not the low Ca^2^+ requirement for Yop secretion (Julio *et al.*, 2001). Dam methylates the N6 position of adenine in 5’-GATC-3’ sequences. In *Y. enterocolitica,* overproduction of Dam alters motility, invasion, and results in increased amounts of rough lipopolysaccharide (LPS) lacking O-antigen side chains (Fälker *et al.*, 2007). Together, these results suggest that Dam sites may show variation in DNA methylation in different environmental conditions including temperature.

Until recently, high-throughput analysis of genomic epigenetic markers was limited to analyzing 5mC via bisulfite conversion followed by sequencing, such as Sanger, pyrosequencing, or whole genome amplification (Tost & Gut, 2007; Zilberman & Henikoff, 2007). The limitation was significant since many studies indicate bacterial DNA methylation centers on N6mA as the modified base (Marinus & Casadesús, 2009; Reisenauer *et al.*, 1999; Wion & Casadesús, 2006). With the advent of single molecule real-time (SMRT) sequencing, a method for sequencing the methylome of bacteria to identify site modifications of adenine exists (Davis *et al.*, 2013; Fang *et al.*, 2012; Murray *et al.*, 2012). SMRT sequencing allows analysis of the native prepared DNA without the requirement of a whole genome amplification, therefore revealing the DNA as the organism has modified it *in vivo* (Flusberg *et al.*, 2010).

In this study, we tested the hypothesis that the *Y. enterocolitica* genome has temperature-dependent differences in DNA methylation. We compared the methylome of *Y. enterocolitica* DNA isolated from cells grown at 22°C and 37°C, providing the first systematic analysis of the epigenetic modifications in this important pathogen. We identified Dam sites with disparate methylation patterns dependent upon growth temperature, and characterized them in relation to promoter regions or coding sequences.

## METHODS

### Bacterial strains, culture media, and DNA techniques

*Y. enterocolitica* strain 8081v (R^-^M+) (Kinder *et al.*, 1993) was grown to late exponential phase in Luria-Bertani broth at either 22°C or 37°C. The presence of the virulence plasmid (pYV) was verified both by PCR and plating on Congo Red LB plates in calcium-chelating conditions (CR-MOX). Genomic DNA was isolated using Sigma (St. Louis) GenElute Bacterial Genomic DNA Kit according to the manufacturer’s protocol.

### SMRT sequencing

SMRT sequencing analyzes the rate at which nucleotides are incorporated on the complimentary replicating strand and compares these kinetics to an *in silico* control. The resulting inter-pulse duration (IPD ratio) provides a kinetic profile that identifies modified bases (Schadt *et al.*, 2013). Libraries of *Y. enterocolitica* 8081v cultures grown at each temperature (22°C and 37°C) were prepared for SMRT sequencing via circular consensus sequencing using a library construction protocol described previously (Travers *et al.*, 2010). Briefly, native chromosomal and plasmid DNA preparations were sheared to an average size of 500bp via adaptive focus acoustics (Corvaris, Woburn, MA, USA), end-repaired, and A-tailed and hairpin adapters with a single T-overhang were ligated. Primers were annealed and sequenced on the Pacific Biosciences RS instrument using C2 chemistry. These libraries were sequenced to a mean coverage depth of 228 (22°C) and 178 (37°C) across the chromosome, and 149 and 68 respectively across the virulence plasmid; average read length was 2656 and 2682, respectively. Reads were mapped to the reference genome (RefSeq NC_008791 and NC_008800 for the chromosome and virulence plasmid, respectively) using BLASR (Chaisson & Tesler, 2012). Base modification and motif detection were performed using the Modification and Motif Detection protocol in the software program SMRTPipe v.1.3.3. Positions with coverage of >25 and Score (QV) of >40 were considered modified. The Score (QV) equals −10*log(p-value), where the p-value was determined from a t-test between the sample and the *in silico* control derived from whole genome amplified (WGA) (i.e. unmethylated) samples (http://pacb.com/applications/base_modification/index.html). Due to the lower read depth coverage on the virulence plasmid, sites of adenine methylation with coverage >10 and an IPD ratio above 3 were considered modified, and was further confirmed because each point of modification was located at known methylation sites.

## RESULTS

### DNA sequence analysis confirmed four genes encoding potential methyltransferase enzymes

The NEB REBASE database identifies 4 potential methyltransferase enzymes (Roberts *et al.*, 2009) which were confirmed by the genomic sequencing of *Y. enterocolitica* strain 8081. The previously reported restriction modification (R-M) *yenI* system (Kinder *et al.*, 1993) was present at loci YE1808. The *yenI* operon encoded a restriction enzyme and a methyltransferase. The YenI restriction enzyme is PstI-like, recognizing the sequence 5’-CTGCAG-3’. Three ‘orphan’ methyltransferase enzymes, i.e., lacking a corresponding restriction enzyme, were also identified: YE3972 and YE2361 encoded DNA adenine methyltransferases (Dam) that recognize 5’-GATC-3’. Previous studies show phenotypic changes following *dam* overexpression in *Yersinia* involve overexpression of loci YE3972 (Fälker *et al.*, 2005; 2006; 2007). A third ‘orphan’ methylase identified as YE2362, was a putative DNA cytosine methyltransferase (Dcm). The coding sequence for this putative Dcm overlapped the 3’-end of the YE2361 Dam methylase.

### Inter-pulse duration analysis identified methylated nucleotides

Our sequencing had increased read-depth coverage of DNA isolated from cells grown at 22°C (222) compared to the sequencing read-depth coverage from cells grown at 37°C (178). This difference is depicted in Figure 2 A and B. The software package to analyze DNA genomic sequences by Pacific Biosciences includes a MEME-chip analysis to identify DNA base modifications. A benefit of SMRT sequencing is that two data types exist in which to determine these base modifications. A quality value (QV) score can be determined by calculating −10*log (p-value) compared to a whole genome amplified (unmethylated) *in silico* control. However, QV scoring has a strong dependence on sequence coverage (number of reads over a given position), so reliability of this value increases as coverage increases. Base modifications can also be determined using the inter-pulse duration (IPD) ratio when sequenced samples differ in their coverage. Comparing the IPD ratio at each base across the genome using this method alleviated bias due to sequence coverage differences (Fig. 3). Therefore, we used the IPD ratio, vs QV scoring, as our kinetic metric to compare DNA prepared from cells grown at each temperature.

Throughout, the following symbols depict DNA methylation states: ■■-denotes fully methylated sites; □■-denotes hemimethylated sites; and □□-denotes unmethylated sites. In addition, patterns that varied with temperature are depicted as methylation condition separated by an arrow indicating the shift from 22°C to 37°C, e.g. □□→□■.

### *Y. enterocolitica* DNA had no evidence of Dcm (5mC) methylation but complete methylation of all *YenI* restriction sites at both 22°C and 37°C

There was no evidence of N4mC or 5mC base modification (Figs. 2 and 3), despite adequate sequence read-depth coverage. Therefore, we concluded that the Dcm methylase (YE2362) was not active in the LB-growth conditions at the two temperatures tested. In contrast, IPD analysis of all 565 YenI (5’-CTGCAG-3’) restriction endonuclease sites showed each adenine was methylated (N6mA) at both temperatures on both DNA strands (Table 1). This complete methylation was expected because these sequences are targets for DNA restriction. These sites are present throughout the genome so identification of adenine methylation at these sites served as an internal control for identifying N6mA at non-YenI sites. The genes and their chromosome positions are depicted in Fig. 1.

**Figure 1.**
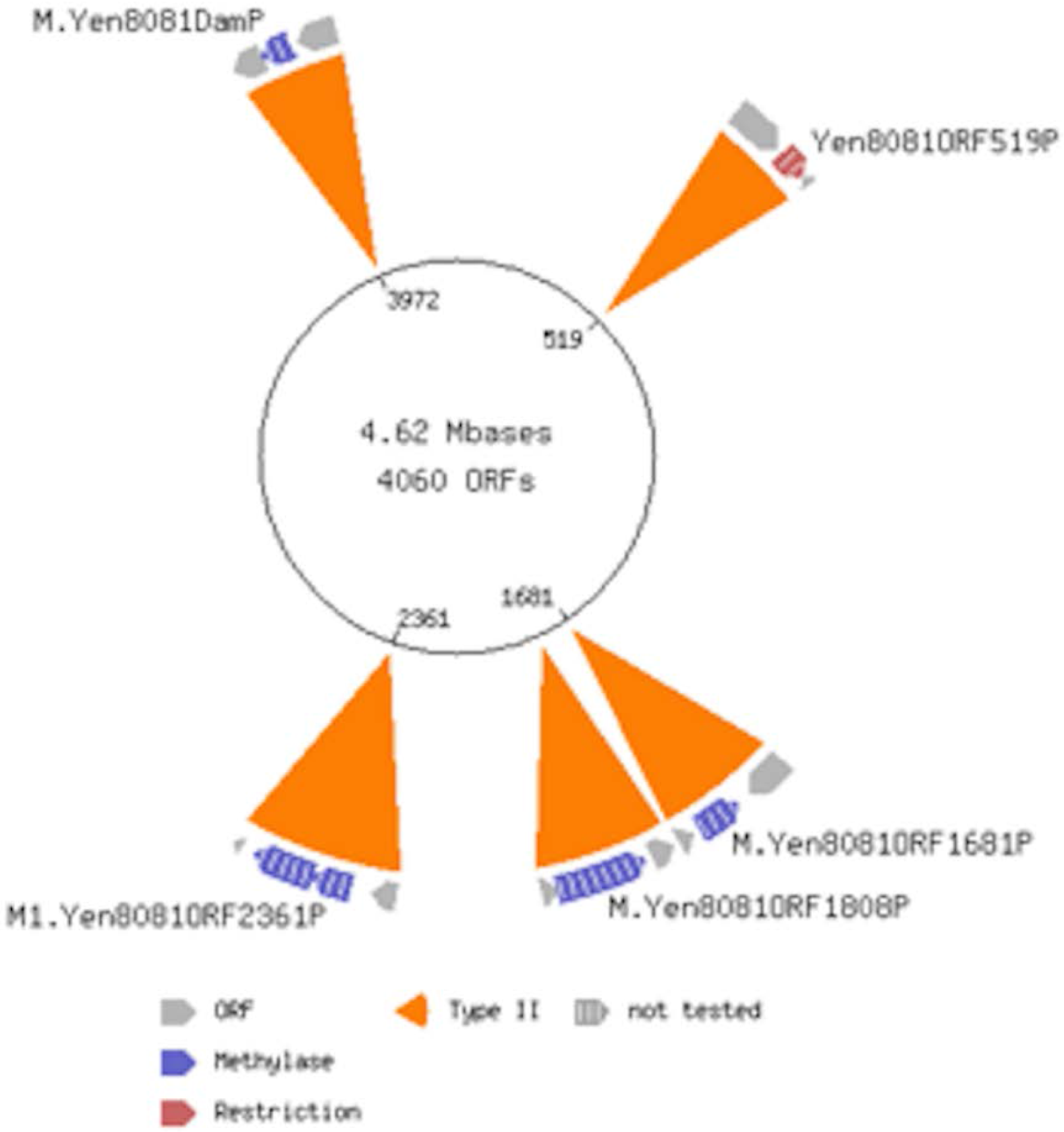
*Y. enterocolitica* sequence analysis identifies 4 methyltransferases. The figure was generated by the REBASE program provided by New England Biolabs. YE0519 is a putative restriction endonuclease with unidentified motif specificity. YE1681 is a putative orphan methyltransferase with unidentified motif specificity, ‘orphan’ referring to the absence of a corresponding restriction endonuclease. YE1808, encodes the YenI restriction modification system recognizing motif 5’-CTGCAG-3’ previously described (Thomson *et al.*, 2006). YE2361 is identified as a DNA adenine methyltransferase (Dam) recognizing 5’-GATC-3’. This ORF overlaps by 4bp with the downstream ORF, YE2362, which, according to NCBI, encodes a DNA cytosine methyltransferase (Dcm). YE3972 is an orphan Dam having recognition specificity to 5’-GATC-3’ as reported previously (Fälker *et al.*, 2005; 2006; 2007).

**Figure 2.**
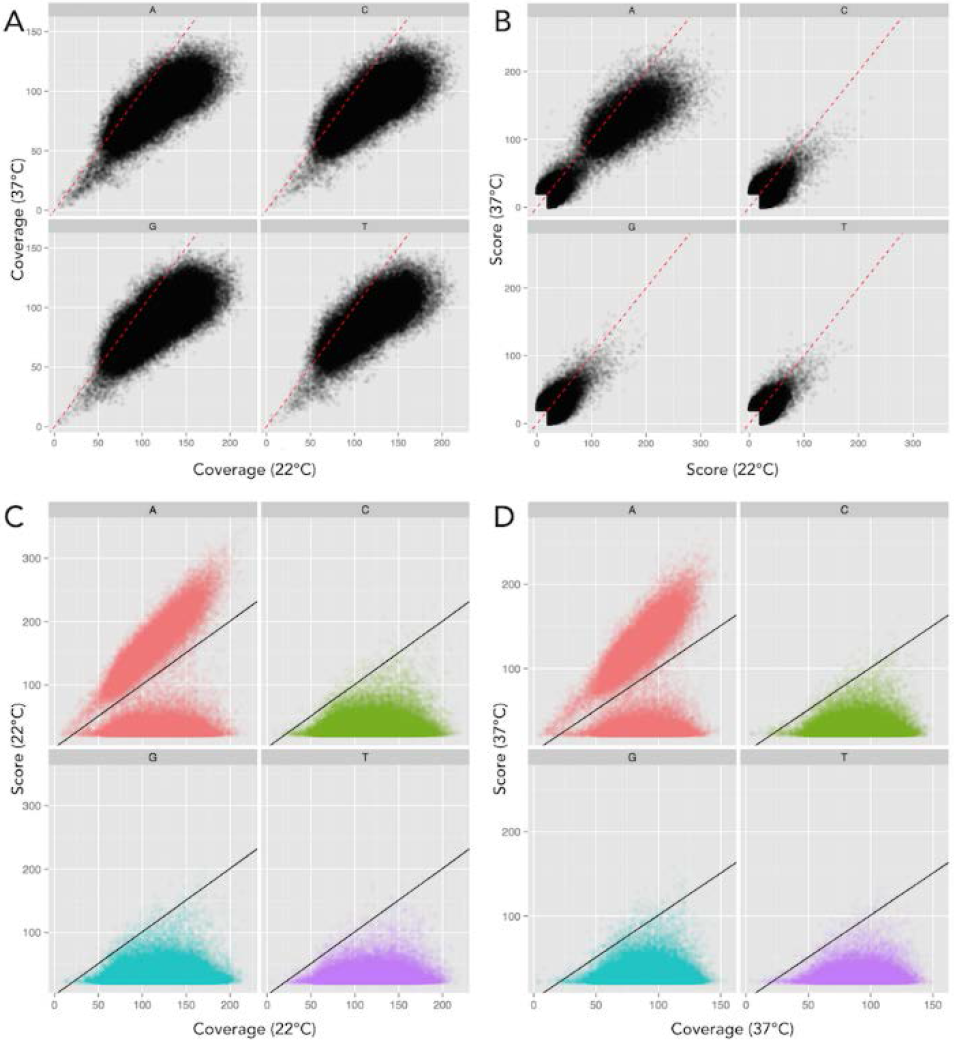
Base modification score to detect methylation is influenced by coverage bias. DNA methylation can be determined by an increased Quality Value (QV) score (-log*10(P-value) compared to unmodified *in silico* control) (Murray *et al.*, 2012). However, the scatter plot in panel (A) illustrates the coverage of the 37°C sample (x-axis) is significantly higher than that of the 22°C sample (y-axis), shown with a 1:1 correlation in the dotted red line. QV score has a strong dependence on coverage, increasing as coverage increases, thus Panel (B) reveals a similar trend when comparing scores as in Panel A, with scores skewing toward the 37°C sample. Therefore, this illustrates that QV scores alone provides a poor metric for comparison between each sample. (C,D) Scatter plot of sequencing coverage and QV score for all genomic positions at 22°C and 37°C respectively, again separated for each base. The threshold for detected base modification above background is indicated by the black line. Scatter plot distributions show significantly higher scores for many adenine (red) residues above the background, whereas no increased scores are evident for all other bases, thus indicating only adenine residues are methylated.

**Figure 3.**
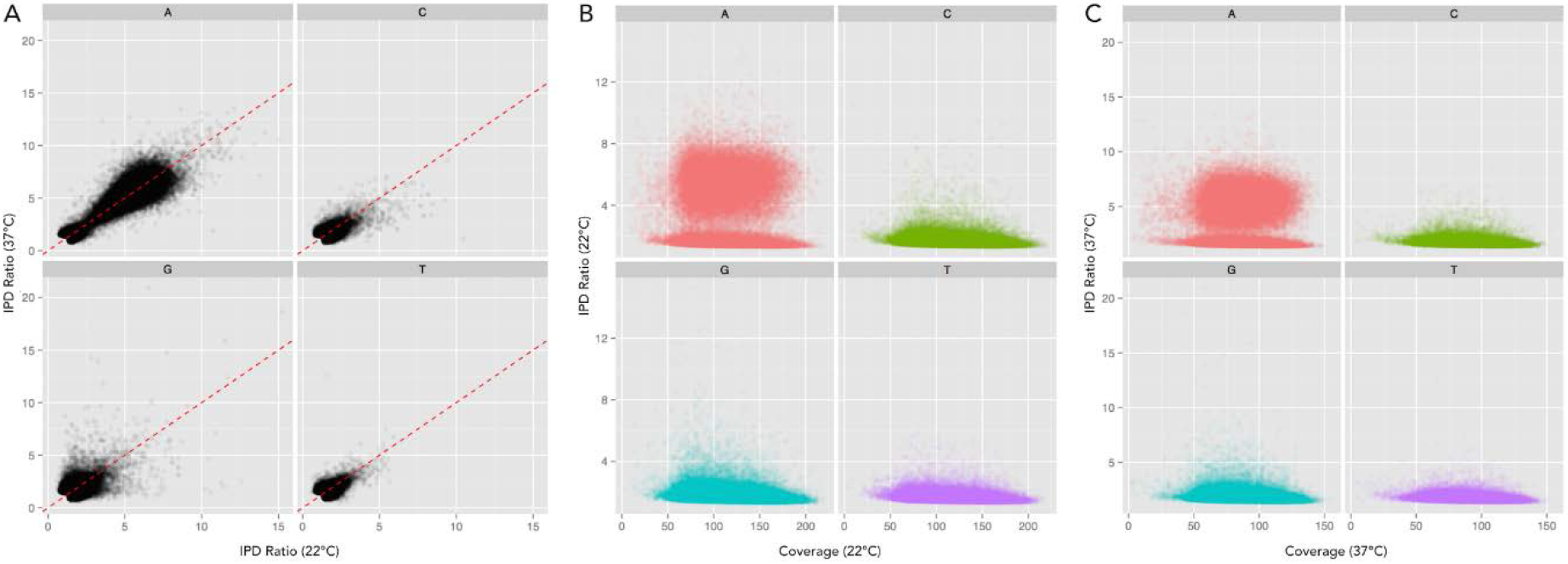
Interpulse duration (IPD) ratio provides an unbiased means of comparing samples with different coverage. (A) Scatter plot comparison of the IPD ratios for all residues from each sample, 22°C on x-axis and 37°C on y-axis, showing no bias toward either sample (1:1 correlation in dotted red line). This permits an unbiased comparison of each sample, indiscriminate of coverage, thus providing a good metric for comparison of each sample. (B,C) Scatter plot of sequencing coverage and IPD ratio for all genomic positions at 22°C and 37°C respectively, separated for each base. Increased IPD ratio above baseline indicates DNA methylation. In each sample, many adenine (red) residues show an increased IPD ratio, revealing adenine methylation.

**Table 1.** Comparison of *YenI*, Dam, and Dcm methylation sites at 22°C and 37°C.

| MOTIF | TOTAL #<br>of Motifs | 22° |  | 37° |  |
| --- | --- | --- | --- | --- | --- |
|  |  | # of<br>Atypical<br>Methylation<br>Detected | # of Fully<br>Methylation<br>Detected | # of<br>Atypical<br>Methylation<br>Detected | # of Fully<br>Methylation<br>Detected |
| 5'-GATC-3' (Dam) |  |  |  |  |  |
| Chromosome | 26664 | 28 | 26636 | 50 | 26614 |
| pYV Plasmid | 454 | 7 | 447 | 7 | 447 |
| 5'-CTGCAG-3' (YenI) |  |  |  |  |  |
| Chromosome | 554 | 0 | 554 | 0 | 554 |
| pYV Plasmid | 11 | 0 | 11 | 0 | 11 |
| 5'-CCWGG-3' (Dcm) |  |  |  |  |  |
| Chromosome | 7237 | 0 | 0 | 0 | 0 |
| pYV Plasmid | 88 | 0 | 0 | 0 | 0 |

<sup>a</sup>Methylated base is underlined.

### *Y. enterocolitica* DNA had different Dam methylation patterns at 22°C and 37°C

Genomic DNA sequence analysis identified 27,118 5’-GATC-3’ Dam methylase sites in the *Y. enterocolitica* genome. The chromosome contained 26,664 sites and the pYV (virulence plasmid) contained 454 sites. IPD analysis showed that the majority of these Dam sites were fully methylated on both DNA strands at both temperatures analyzed (Table 1). Importantly, a subset of 42 Dam sites had different temperature-dependent methylation patterns (Table 2). In addition to these 42 sites, 17 Dam sites remained unmethylated at both temperatures (□□→□□, Table 3). Temperature-dependent Dam methylation patterns were found throughout the genome in both regulatory and gene open reading frame sequences. The genes identified were classified as coding for, putative regulators, ribosomal RNAs, membrane-associated proteins, metabolic proteins, virulence proteins, or for hypothetical proteins of no known function. We found no evidence of cytosine methylation in DNA from cells grown in these experimental conditions. Finally, all YenI restriction endonuclease sites were fully methylated on both DNA strands at both temperatures. The specifics of the differential methylation are in Tables 2 and 3 and outlined in the following paragraphs.

**Table 2.** Dam sites showing temperature-dependent methylation differences at 22°C and 37°C.

| Position | 22° a | 37° a | Name | Location b | Cat. c | Description |
| --- | --- | --- | --- | --- | --- | --- |
| 20357 | ■ ■ | □ □ | YEr002 | ORF | rR | 23S rRNA |
| 315457 | ■ ■ | ■ □ | YEr007 | ORF, -160 from 3' end | rR | 16S rRNA |
| 315631 | ■ □ | ■ □ |  | +16 from 3' end of 16S rRNA |  |  |
| 356036 | ■ ■ | ■ □ | YEr010 | ORF, -158 from 3' end | rR | 16S rRNA |
| 356210 | ■ ■ | ■ □ |  | +16 from 3' end of 16S rRNA |  |  |
| 373662 | □ □ | ■ □ | YE0315 | -172 from TSS | M | putative membrane transport proteins, similar to Salmonella TonB |
|  |  |  | YE0316 | -206 from TSS | MT | putative DNA-binding protein |
| 399373 | ■ ■ | □ □ | YE0335 | -75 from TSS | MT | HemN, coproporphyrinogen III oxidase |
| 623091 | □ □ | ■ □ | YE0546 | ORF | M | putative glycosyltransferase |
| 888993 | □ □ | ■ □ | YE0762 | ORF | MT | CysD, sulfate adenylyltransferase subunit 2 |
| 1042835 | □ □ | ■ ■ | YE0914 | -36 from TSS | R | putative LysR-type transcriptional regulator |
| 1104202 | □ □ | ■ □ | YE0981 | -74 from TSS | H | hypothetical protein |
| 1111932 | ■ ■ | ■ □ | YE0990 | ORF | MT | AYP/GTP-binding protein |
| 1113753 | ■ ■ | ■ □ | YE0992 | ORF | H | hypothetical protein |
| 1228189 | □ □ | ■ □ | YE1098 | -182 from TSS | MT | GutA, pts system, glucitol/sorbitol-specific iic2 component |
| 1403906 | □ □ | ■ □ | YE1259 | -51 from TSS | R | putative PadR-like family transcriptional regulator |
| 1468949 | ■ ■ | ■ □ | YE1322 | ORF | V | putative RTX-family protein |
| 1469615 | ■ ■ | ■ □ |  |  |  |  |
| 1471616 | ■ ■ | □ □ |  |  |  |  |
| 1471736 | ■ ■ | ■ □ |  |  |  |  |
| 1471902 | ■ ■ | ■ □ |  |  |  |  |
| 1879399 | ■ ■ | ■ □ | YE1683 | ORF | R | putative prophage encoded two-component system response regulator |
| 1881251 | ■ ■ | ■ □ | YE1684 | ORF | R | putative prophage encoded two-component system histidine kinase |
| 2354334 | ■ ■ | ■ □ | YE2154 | ORF. near 5' | MT | Rnt, ribonuclease T |
| 2436018 | ■ ■ | ■ □ | YE2225 | -288 from TSS | R | putative cyclic-di-GMP phosphodiesterase |
| 2598581 | ■ ■ | ■ □ | YE2407 | ORF | V | putative hemolysin |
| 2602469 | ■ ■ | ■ □ | YE2409 | +30 from 3' end of mviN | MT | MviN, putative membrane-associated protein |
| 2852693 | ■ ■ | ■ □ | YE2635 | ORF |  | putative metallo-beta-lactamase superfamily protein |
| 3352496 | ■ ■ | ■ □ | YE3082 | ORF | MT | RfbX, putative O-antigen transporter |
| 3548081 | ■ ■ | ■ □ | YEr017 | ORF | rR | 23S rRNA |
| 3549372 | ■ □ | □ □ |  |  |  |  |
| 3550740 | ■ ■ | ■ □ | YEr018 | +17 from 3' end of 16S rRNA | rR | 16S rRNA |
| 3655022 | ■ ■ | ■ □ | YE3343 | ORF. near 3' | MT | OutL, general secretion pathway protein L |
| 3739389 | □ □ | ■ □ | YE3423 | -85 from TSS | R | ArsR-family transcriptional regulator |
|  |  |  | YE3424 | -66 from TSS | M | putative zinc metallopeptidases |
| 4243325 | ■ ■ | □ □ | YEr022 | +17 from 3' end of 16S rRNA | rR | 16S rRNA |
| 4441739 | □ □ | ■ ■ | YE4070 | -100 from TSS | MT | putative oligogalacturonate-specific porin protein |
| 2443 | ■ ■ | ■ □ | YEP0004 | ORF | V | YopQ, virulence plasmid protein |
| 2654 | ■ □ | ■ □ |  | ORF |  |  |
| 15967 | ■ □ | ■ ■ | YEP0018 | ORF | MT | LcrD, low calcium response locus membrane protein d |
| 21305 | ■ □ | ■ □ | YEP0026 | ORF | V | YscP, putative type III secretion protein |
| 23409 | ■ □ | ■ ■ | YEP0029 | ORF | V | YscS, putative type III secretion protein |
| 24586 | ■ ■ | ■ □ | YEP0031 | ORF | V | YscU, putative type III secretion protein |
| 25258 | ■ □ | ■ ■ |  | ORF |  |  |
| 45942 | ■ ■ | ■ □ | YEP0064 | -86 from TSS | H | putative pseudogene |
| 48481 | ■ □ | ■ ■ | YEP0066 | -158 from TSS | M | putative YadA invasin |
| 66135 | ■ ■ | ■ □ | YEP0096 | ORF | MT | putative plasmid copy control protein |

<sup>a</sup>■ ■ ■, fully methylated; ■ □, hemimethylated; □ □, unmethylated
<sup>b</sup>ORF, Open Reading Frame; TSS, Translational start site.
<sup>c</sup>H, Hypothetical; R, Regulator; rR, ribosomal RNA; M, membrane-associated; MT, metabolic; V, virulence

**Table 3.**
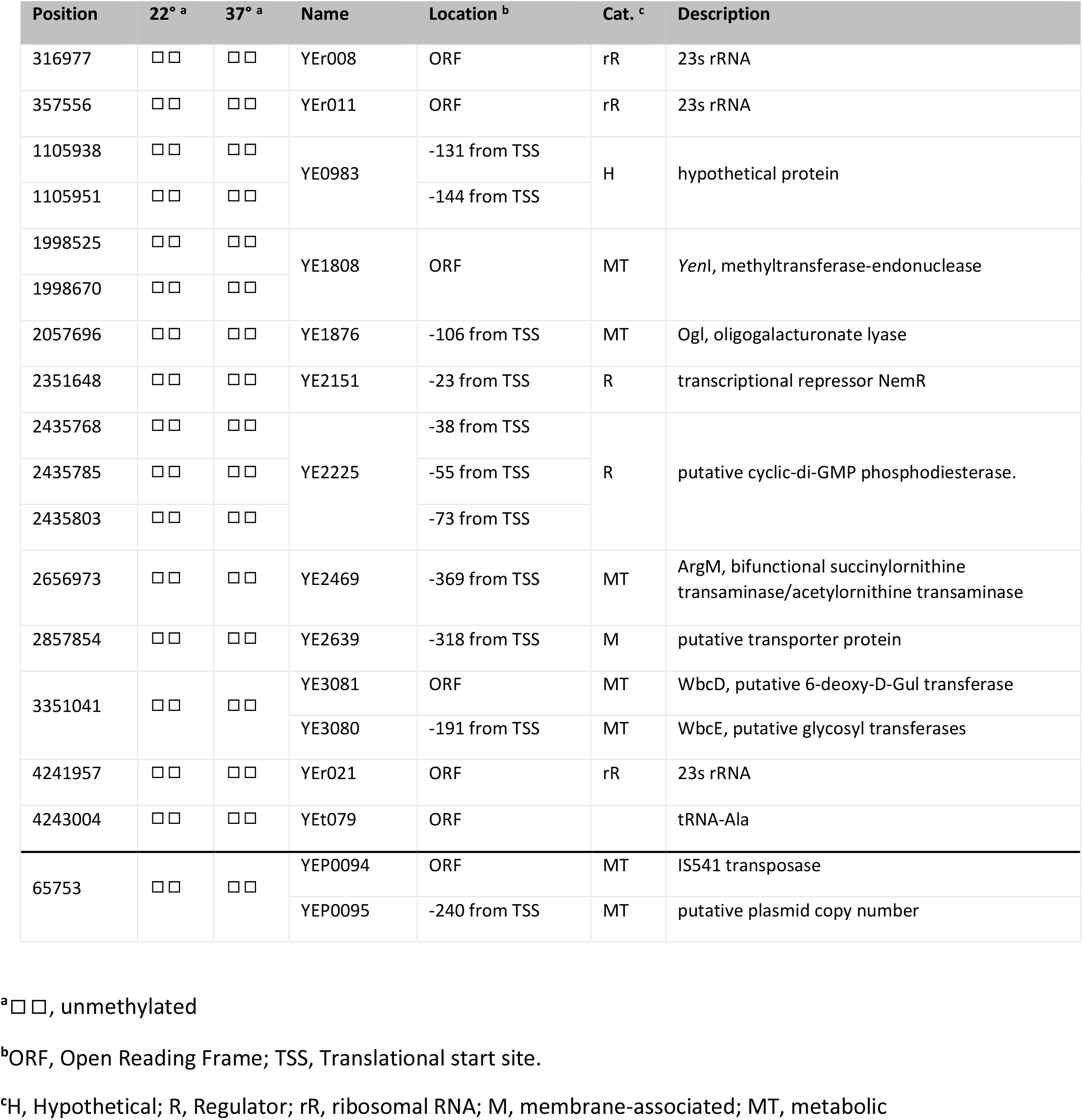
Dam sites unmethylated at both 22°C and 37°C.

| Position | 22°<sup>a</sup> | 37°<sup>a</sup> | Name | Location<sup>b</sup> | Cat.<sup>c</sup> | Description |
| --- | --- | --- | --- | --- | --- | --- |
| 316977 | □□ | □□ | YEr008 | ORF | rR | 23s rRNA |
| 357556 | □□ | □□ | YEr011 | ORF | rR | 23s rRNA |
| 1105938 | □□ | □□ | YE0983 | -131 from TSS | H | hypothetical protein |
| 1105951 | □□ | □□ |  | -144 from TSS |  |  |
| 1998525 | □□ | □□ | YE1808 | ORF | MT | *YenI*, methyltransferase-endonuclease |
| 1998670 | □□ | □□ |  |  |  |  |
| 2057696 | □□ | □□ | YE1876 | -106 from TSS | MT | Ogl, oligogalacturonate lyase |
| 2351648 | □□ | □□ | YE2151 | -23 from TSS | R | transcriptional repressor NemR |
| 2435768 | □□ | □□ | YE2225 | -38 from TSS | R | putative cyclic-di-GMP phosphodiesterase. |
| 2435785 | □□ | □□ |  | -55 from TSS |  |  |
| 2435803 | □□ | □□ |  | -73 from TSS |  |  |
| 2656973 | □□ | □□ | YE2469 | -369 from TSS | MT | ArgM, bifunctional succinylornithine transaminase/acetylornithine transaminase |
| 2857854 | □□ | □□ | YE2639 | -318 from TSS | M | putative transporter protein |
| 3351041 | □□ | □□ | YE3081 | ORF | MT | WbcD, putative 6-deoxy-D-Gul transferase |
|  |  |  | YE3080 | -191 from TSS | MT | WbcE, putative glycosyl transferases |
| 4241957 | □□ | □□ | YEr021 | ORF | rR | 23s rRNA |
| 4243004 | □□ | □□ | YEt079 | ORF |  | tRNA-Ala |
| 65753 | □□ | □□ | YEP0094 | ORF | MT | IS541 transposase |
|  |  |  | YEP0095 | -240 from TSS | MT | putative plasmid copy number |

<sup>a</sup>□□, unmethylated
<sup>b</sup>ORF, Open Reading Frame; TSS, Translational start site.
<sup>c</sup>H, Hypothetical; R, Regulator; rR, ribosomal RNA; M, membrane-associated; MT, metabolic

Two Dam sites were unmethylated at 22°C and methylated at 37°C (□□→■■) localized in the 5’-regulatory regions of YE0914 and YE4070, at 36 and 100 bp respectively 5’ from the predicted AUG start codons (Table 2). YE0914 is identified by NCBI as a hypothetical protein but EMBL’s Pfam sequence analysis identified it as a LysR-type transcription regulator (LTTR) (Finn *et al.*, 2016). LTTRs contain a helix-turn-helix DNA-binding domain and are among the most abundant type of transcriptional regulators within prokaryotes (Henikoff *et al.*, 1988; Maddocks & Oyston, 2008). YE4070 codes for a putative outer membrane oligogalacturonate-specific porin, KdgM.

Four Dam sites were fully methylated at 22°C and unmethylated at 37°C (■■→□□). These included the YE0335 *(hemN)* gene coding for coproporphyrinogen III oxidase with the Dam site 75 bp 5’ from the start codon. Two (■■→□□) sites were within ribosomal RNA genes, both a 16s rRNA and 23s rRNA, and one was within the coding region of YE1322, a putative RTX-family protein, associated with type I secretion system pore-forming toxins. YE1322, with a predicted open reading frame of 6333 bp, contains 17 additional Dam sites, four of which had full methylation at 22°C and hemimethylation at 37°C (■■→■□).

Of the seven Dam sites unmethylated at 22°C and hemimethylated at 37°C (□□→□■), five were in probable regulatory regions and two were within gene coding regions. Of the five Dam sites in intergenic regulatory regions, three were in genes identified as possible regulatory proteins. The first regulatory gene identified with this category was YE1259, with a Dam site 51 bp 5’ from the AUG start codon. YE1259 was identified as a PadR-like transcriptional regulator. The second regulatory gene in this category was YE3423, with a Dam site 85 bp 5’ from the AUG start codon. YE3423 codes for an ArsR-family transcriptional regulator. Of note, this Dam sites is positioned between two divergently-transcribed genes, placing it 66 bp 5’ from the AUG start codon of YE3424. YE3424 codes for a putative zinc metallopeptidase. A BLAST search of the YE3424 predicted amino acid sequence had 70% identity and 83% similarity to enterohemorrhagic *E. coli* (EHEC) metallopeptidase, SprT, a type III secretion system effector. The third regulator gene in this category of Dam methylation (□□→□■) was YE0316 with a Dam site 206 bp 5’ from the AUG start codon. YE0316 codes for putative DNA-binding protein with 53% similarity to sigma-70 (*fecI*). Of note, this Dam sites is also positioned 172 bp 5’ from the AUG start codon of divergently transcribed YE0315. YE0315 codes for a membrane transport protein with high similarity to *Salmonella typhimurium* TonB. Two additional genes showed this pattern of methylation (□□→□■): YE0981, encoding a hypothetical protein with a Dam site 74 bp 5’ from the start codon; and YE1098 with the Dam site 51 bp 5’ from the start codon. This gene encodes GutA, also referred to as SrlA, a glucitol/sorbitol-specific IIC2 component, a subunit of the phoshotransferase system (Meadow *et al*., 1990).

Of the 24 Dam sites fully methylated at 22°C and hemi-methylated at 37°C (■■→■□), 22 were located within structural gene coding sequences (Table 2). Of the two sites in regulatory regions, one ■■→■□ Dam site is positioned 288 bp 5’ from the AUG start codon of YE2225, a predicted cyclicdi-GMP phosphodiesterase with a conserved EAL domain. This enzyme inactivates cyclic-di-GMP, a common bacterial secondary messenger (Romling *et al.*, 2013). Interestingly, within the promoter of YE2225, three additional Dam sites had differential methylation patterns. The three Dam sites most proximal to the coding region, 38, 55, and 73 bp 5’ from the AUG start codon, remained unmethylated at each temperature. It is noteworthy that this regulatory region, from 38 to 288 5’ from the start AUG codon, contains four Dam sites that show atypical methylation patterns (Tables 2 and 3). Statistically, only one Dam site was predicted over a span of 256 bp. The second Dam site in a potential regulatory region is on the virulence plasmid, 86 bp from the 5’ start of YEP0064, a putative pseudogene. This pattern of methylation (■■→■□) was also prominent in rRNA genes. We identified five Dam sites in 16S and 23S rRNA genes, which were position-specific at the 3’-ends. This included four of the seven 16S ribosomal RNA genes (YEr007, YEr010, YEr018, and YEr022) with this conserved pattern of atypical methylation (Tables 2 and 3).

Seventeen Dam sites were identified to be unmethylated at each temperature (□□→□□). Ten were located in probable regulatory regions (Table 3). As described above, three unmethylated Dam sites were located 5’ to the start of YE2225, the putative cyclic-di-GMP phosphodiesterases described above. Two were at 131- and 144 bp 5’ from the start codon of YE0983, a hypothetical protein. The remaining unmethylated Dam sites were 5’ to the AUG start codon of an oligogalacturonate lyase, a bifunctional transaminase, a putative transporter, a putative glycosyl transferase, and YE2151, a TetR-like transcriptional regulator similar to NemR. (Table 3).

## DISCUSSION

The most significant finding of this work identified two subsets of Dam methylation sites with temperature-dependent Dam methylation patterns or Dam sites that remained unmethylated regardless of temperature. To our knowledge, this is the first systematic analysis of temperature-dependent DNA methylation of a facultative bacterial pathogen. Importantly, IPD ratio, rather than the traditional QV scoring, was the kinetic metric used to compare DNA so that DNA sequence coverage differences between cells grown a different temperatures was not an issue. Both categories of Dam sites were localized either within regulatory regions (5’ from the predicted AUG start site) or within gene open-reading frames. Among the genes with differential methylation were those encoding regulatory, virulence, metabolic, membrane-associated proteins, and both 23s and 16s rRNA genes. Importantly, we found no evidence of Dcm methylation at predicted Dcm sites (5’-CCWGG-3’) on the chromosome nor on the pYV. To determine if the temperature-dependent methylation patterns identified correlated with gene expression we capitalized on the *Y. enterocolitica* comprehensive transcriptome analyzes conducted by Bent *et. al.* (Bent *et al.*, 2015). This study is a comprehensive RNA-seq analysis of *Y. enterocolitica* 8081v transcripts prepared from cultures grown at 25°C in LB broth and from 37°C cultures grown in (i) conditioned RPMI, (ii) in surface contact with mouse macrophages, or (iii) internalized by mouse macrophages.

The LTTR at loci YE0914 has a Dam site in the predicted regulatory region that is unmethylated at 22°C and fully methylated at 37°C (□□→■■) exemplifying a gene that would require two rounds of DNA replication to restore the unmethylated state after a temperature downshift. This gene is highly conserved among the enteric bacteria with 88% identity to an LTTR in *Serratia marscesens* and 76% identity to an LTTR in *Shigella.* To our knowledge, YE0914, nor its enteric homologs, have been characterized. LTTRs are among the most abundant types of transcriptional regulators present in bacteria, and control a diverse subset of genes, including motility, metabolism, and virulence (Maddocks & Oyston, 2008). The *E. coli* LTTR, OxyR, activates one of its many targets, the phage Mu *mom* gene, only when three Dam sites upstream of the promoter are methylated (Bölker & Kahnmann, 1989). Therefore, there is precedence for LTTR expression correlating with promoter methylation. Transcription of YE0914 in the RNA-seq transcriptomic analysis by Bent *et al.* (Bent *et al.*, 2015) does not show temperature-dependent regulation between *Y. enterocolitica* cells grown in LB at 26°C compared to conditioned RPMI at 37°C (Bent *et al.*, 2015). However, the p-value in this study for YE0914 at these two temperatures is very high (0.65-0.93), suggesting significant heterogeneity in expression within sampled populations. Heterogenic expression is also indicative of epigenetic mechanisms whereby bacteria modify gene expression to “bet hedge” within otherwise clonal populations (Casadesús & Low, 2013).

Among the regulatory genes identified with unmethylated Dam site(s) (□□→□□) upstream of the translational start was the TetR-like transcriptional regulator YE2151. This gene has a predicted 64% amino acid similarity (52% identity) to *E. coli* NemR (Gray *et al.*, 2013). NemR activates several stress-response genes required for survival when cells are exposed to reactive oxygen species such as hypochlorus acid (Hassett & Cohen, 1989). The transcriptomic analysis by Bent *et al.* (Bent *et al.*, 2015) shows significantly increased expression of YE2151 at elevated temperature. However, based on our methylation analysis, methylation did not appear to play a role in regulation since the unmethylated state of the Dam site identified did not change with temperature. It is noteworthy that this stress-response gene is localized near the terminus of DNA replication. Genes expressed during stress or stationary phase are positionally-biased to be near the terminus and show a requirement for reduced supercoiling for expression (Muskhelishvili & Travers, 2013; Travers & Muskhelishvili, 2015). Previous studies by our laboratory have likewise demonstrated that temperature regulation in *Y. enterocolitica* correlates to changes in DNA supercoiling. DNA methylation can affect histone-like protein binding which in turn dictate regional DNA domain confirmation. As such, lack of methylation at the chromosome terminus may influence gene expression indirectly through Dam site protection.

Also showing differential regulation in its upstream regulatory region, was the regulatory gene, YE1259 (□□→■□), coding for a PadR-like transcriptional regulator. PadR family transcriptional regulators are involved in *Vibrio* virulence. For example, AphA is involved in the regulation of motility and virulence in *Vibrio parahaemolyticus* (Wang *et al.*, 2013). In *V. parahaemolyticus,* both an LTTR (AphB) and AphA co-regulate acetoin production and motility (Kovacikova *et al.*, 2004; 2005). Notably these phenotypes are temperature regulated in *Y. enterocolitica,* where acetoin and flagella are only produced at low temperature (Chester & Stotzky, 1976). The transcriptomic analysis by Bent *et. al.* (Bent *et al.*, 2015) however, reveals little variation in temperature expression of YE1259. YE1098, coding for GutA, also showed the same pattern of methylation as YE1259 (□□→■□). van der Woude *et al.* (van der Woude *et al.*, 1998) reported that the Dam site 44 bp from the *E. coli gutABD* transcription start site is unmethylated. GutR and CRP compete for binding at this Dam site, and GutR binding prevents methylation. However, changing the Dam site in *E. coli* did not alter expression, suggesting that methylation does not play a direct role in *gutABD* regulation. In *Y. enterocolitica* we noted the presence of a putative CRP-binding site overlapping this Dam site suggesting similar regulation. The fact that we saw a transition in methylation pattern with growth temperature suggests methylation of this operon may be sensitive to other environmental conditions. Indeed, Bent *et al.* (Bent *et al.*, 2015) show significant activation of *gutA* at 37°C.

The most extensive atypical Dam methylation was seen in YE2225. Over a span of 288 bp upstream of the translational start site are four Dam sites, three of which remained unmethylated at both temperatures and one, the most distal from the AUG start site, went from fully methylated at 25°C to hemimethylated at 37°C (■■→■□). YE2225 encodes a putative cyclic-di-GMP phosphodiesterase (PDE) with a conserved EAL domain required for hydrolysis of the common bacterial signaling messenger cyclic-di-GMP (Romling *et al.*, 2013). Transcriptomic analysis by Bent *et al.* (Bent *et al.*, 2015) shows this gene undergoes a steady upregulation when exposed to mouse macrophages grown in RPMI at 37°C reaching a ~5.5-fold peak increase at 60 min. Cyclic-di-GMP plays an essential role during environmental transitions of bacteria. Phenotypes governed by cyclic-di-GMP include virulence gene expression and the transition from the motile to sessile states during biofilm formation (Bobrov *et al.*, 2011). For the related *Y. pestis,* cyclic-di-GMP regulates exopolysaccharide production during biofilm formation in the flea vector. Hydrolysis of cyclic-di-GMP is essential in the transition from flea ambient temperature to mammalian host temperature. Bobrov *et al.* (2011) show mutational inactivation of HmsP, the *Y. pestis* cyclic-di-GMP phosphodiesterase, results in a 1000-fold reduction in virulence. Our results suggest methylation in concert with higher temperature may alter cyclic-di-GMP levels essential for host infection. Interestingly in *V. cholera,* both AphA and AphB repress the activation of the EAL-containing *acgA.* AcgA further shows effects on motility, virulence, and biofilm formation (Kovacikova *et al.*, 2005).

YE0316 (□□→■□) is a homolog of FecI. FecI is a sigma factor which regulates expression of the ferric di-citrate uptake system clustered chromosomally in the iron regulatory region of *E. coli* (Buchanan, 2005). The *fecI* gene is temperature-regulated in *E. coli,* with higher expression at 37°C, and controlled by histone-like nucleoid structuring (H-NS) protein (White-Ziegler & Davis, 2009). This temperature regulation is true for YE0316; expression is increased over eight-fold at 120 mins following 37°C exposure (Bent *et al.*, 2015). Notably, H-NS binding is affected by DNA methylation (White-Ziegler *et al.*, 1998).

Several locations of disparate methylation patterns were within or near ribosomal RNA elements. Of the seven 16S rRNA genes present on the chromosome, four showed temperature variation in methylation. Similarly, of the seven 23S rRNA elements, five showed temperature variation in methylation. The positions for each of these Dam sites were localized to the 3’ end of the coding region. Furthermore, the ribosomal RNAs showing this methylation pattern were localized near the origin of replication. Genomic position does effect expression levels (Gerganova *et al.*, 2015; Muskhelishvili & Travers, 2013).

Finally, we identified 17 Dam sites which remain unmethylated. This number is consistent with unmethylated sites determined for uropathogenic *E. coli* and *C. crescentus* (Blyn *et al.*, 1990; Fang *et al.*, 2012; Kozdon *et al.*, 2013; van der Woude *et al.*, 1998). This suggests these sites are permanently protected by either binding by proteins, local DNA domain structure, or both. It is noteworthy that 14 of these 17 unmethylated sites are clustered in 10 genes positioned near the origin (five sites) or terminus of DNA replication (12 sites).

The lack of cytosine methylation detected in our analysis is consistent with the transcriptome analysis of Bent *et al.* (Bent *et al.*, 2015). Our interpretation of their data indicates either no or minimal expression of YE2361 *(dam)* and YE2362 *(dcm)* at all growth and temperature conditions monitored in their experiments. Conversely, *yenI* (YE1808) and *dam* (YE3972) are expressed constitutively with little temperature variation.

Overproduction of Dam affects motility and invasion abilities in *Y. enterocolitica,* implying that temperature-dependent phenotypic differences may be regulated by temperature-responsive differential methylation (Low *et al.*, 2001). In this study, we showed there is a subset of Dam sites displaying temperature-dependent methylation patterns. Of the Dam sites in regulatory regions, some patterns correlate with temperature regulation when compared to the RNA-seq data of Bent *et al.* (Bent *et al.*, 2015). Surprisingly, we did not see differences in methylation of regulatory regions of key genes involved in TTSS temperature regulation suggesting if methylation is a component of motility and Yop expression, it is indirect. However, the mechanism underlying this regulation by Dam remains unclear, the genes identified here provide targets for further investigation.

## ACKNOWLEDGEMENTS

This work was supported by the University of Idaho Agricultural Experiment Station Hatch projects under award numbers IDA01467 (C.J.H.) and IDA01406 (S.A.M.) and by the National Institutes of Health under award number P20GM103408

## ABBREVIATIONS

SMRT: single-molecule real-time
IPD: interpulse duration
QV: quality value
Dam: DNA adenine methylase
Dcm: DNA cytosine methylase
PDE: phosphodiesterase
LPS: lipopolysaccharide
TLR: Toll-like receptor
TTSS: type-III secretion system
Yops: *Yersinia* outer proteins
N6mA: N6-methyladenine
N4mC: N4-methylcytosine
5mC: 5-methylcytosine
LTTR: LysR-type transcriptional regulator

